# An interaction map of circulating metabolites, immune gene networks and their genetic regulation

**DOI:** 10.1101/089839

**Authors:** Artika P. Nath, Scott C. Ritchie, Sean G. Byars, Liam G. Fearnley, Aki S. Havulinna, Anni Joensuu, Antti J. Kangas, Pasi Soininen, Annika Wennerström, Lili Milani, Andres Metspalu, Satu Männistö, Peter Würtz, Johannes Kettunen, Emma Raitoharju, Mika Kähönen, Markus Juonala, Aarno Palotie, Mika Ala-Korpela, Samuli Ripatti, Terho Lehtimäki, Gad Abraham, Olli Raitakari, Veikko Salomaa, Markus Perola, Michael Inouye

**Author notes:** Correspondence addressed to M.I.

## Abstract

The interaction between metabolism and the immune system plays a central role in many cardiometabolic diseases. We integrated blood transcriptomic, metabolomic, and genomic profiles from two population-based cohorts, including a subset with 7-year follow-up sampling. We identified topologically robust gene networks enriched for diverse immune functions including cytotoxicity, viral response, B cell, platelet, neutrophil, and mast cell/basophil activity. These immune gene modules showed complex patterns of association with 158 circulating metabolites, including lipoprotein subclasses, lipids, fatty acids, amino acids, and CRP. Genome-wide scans for module expression quantitative trait loci (mQTLs) revealed five modules with mQTLs of both *cis* and *trans* effects. The strongest mQTL was in *ARHGEF3* (rs1354034) and affected a module enriched for platelet function. Mast cell/basophil and neutrophil function modules maintained their metabolite associations during 7-year follow-up, while our strongest mQTL in *ARHGEF3* also displayed clear temporal stability. This study provides a detailed map of natural variation at the blood immuno-metabolic interface and its genetic basis, and facilitates subsequent studies to explain inter-individual variation in cardiometabolic disease.

## Introduction

Over the last decade increasing evidence has implicated inflammation as a probable causal factor in metabolic and cardiovascular diseases. Consequently, research has begun to focus on the interplay between immunity and metabolism, or immunometabolism. While it is involved in diverse pathophysiologies, immunometabolism is particularly relevant to diseases of immense global health burden, such as type 2 diabetes (T2D) and atherosclerosis.

For T2D, immune overactivation in adipose tissue has been implicated as a key driver (1,2). Studies have shown that macrophage infiltration and subsequent overexpression of proinflammatory cytokines, such as TNF-a, in adipose tissues is associated with insulin resistance (1,2). Moreover, evidence for metabolic inflammation has been shown in other tissues where, in blood, elevated glucose and free fatty acid levels potentiate IL-1β mediated destruction of pancreatic ß-cells and subsequent T2D progression (3–5). While circulating metabolites are known to be associated with cardiovascular disease (6), inflammation is an increasingly recognised factor in pathogenesis. In atherosclerosis, lipid-induced inflammatory response mechanisms have also been implicated in progression to myocardial infarction (7). In atherogenic lesions, oxidized phospholipids are known to lead to a new macrophage phenotype (8), and cholesterol loading in macrophages promote proinflammatory cytokine secretion (9).

Perhaps surprisingly, few large-scale studies have systematically assessed interactions between the human immune system and metabolites. Recent studies have investigated matched blood transcriptomic and metabolomic profiles to understand their interplay (10–16). However, these studies had modest sample sizes and thus have not had the power to focus on the diverse range of immune processes that interact with circulating metabolites. Furthermore, even fewer have assessed effects of expression quantitative trait locus (eQTL) on immune gene networks. A robust integrated map of immunometabolic relationships and their genetic regulation would provide a foundation for investigating the differential cardiometabolic disease susceptibility amongst individuals while also identifying key target interactions for mechanistic *in vivo* and *in vitro* follow-up.

In this study, we present an integrated immunometabolic map using matched blood metabolomic and transcriptomic profiles from 2,168 individuals from two population-based cohorts. We perform gene coexpression network discovery and cross-cohort replication to identify robust gene modules which encode immune-related functions. Using a high-throughput quantitative NMR metabolomics platform that can separate lipids and lipoprotein sub-fractions as well as quantify a panel of polar metabolites, we identify significant interactions between immune gene modules and circulating metabolites. Genome-wide scans for QTLs affecting immune gene modules identify many *cis* and *trans* loci affecting module expression. Finally, we test the long-term stability of gene modules, their metabolite interactions, and genetic control using a 7-year follow up sampling of 333 individuals.

## Results and Discussion

### Summary of cohorts and data

We analysed genome-wide genotype, whole blood transcriptomic and serum metabolomics data from two population-based cohorts (**Methods** and **Figure 1**). In DILGOM07, 240 males and 278 females aged 25-74 years were recruited (total N = 518). Data was available for a subset of 333 participants from DILGOM07 who were followed up after seven years (DILGOM14). In YFS, relevant data was available for 755 males and 895 females aged 34-49 years (total N = 1,650).

**Figure 1:**
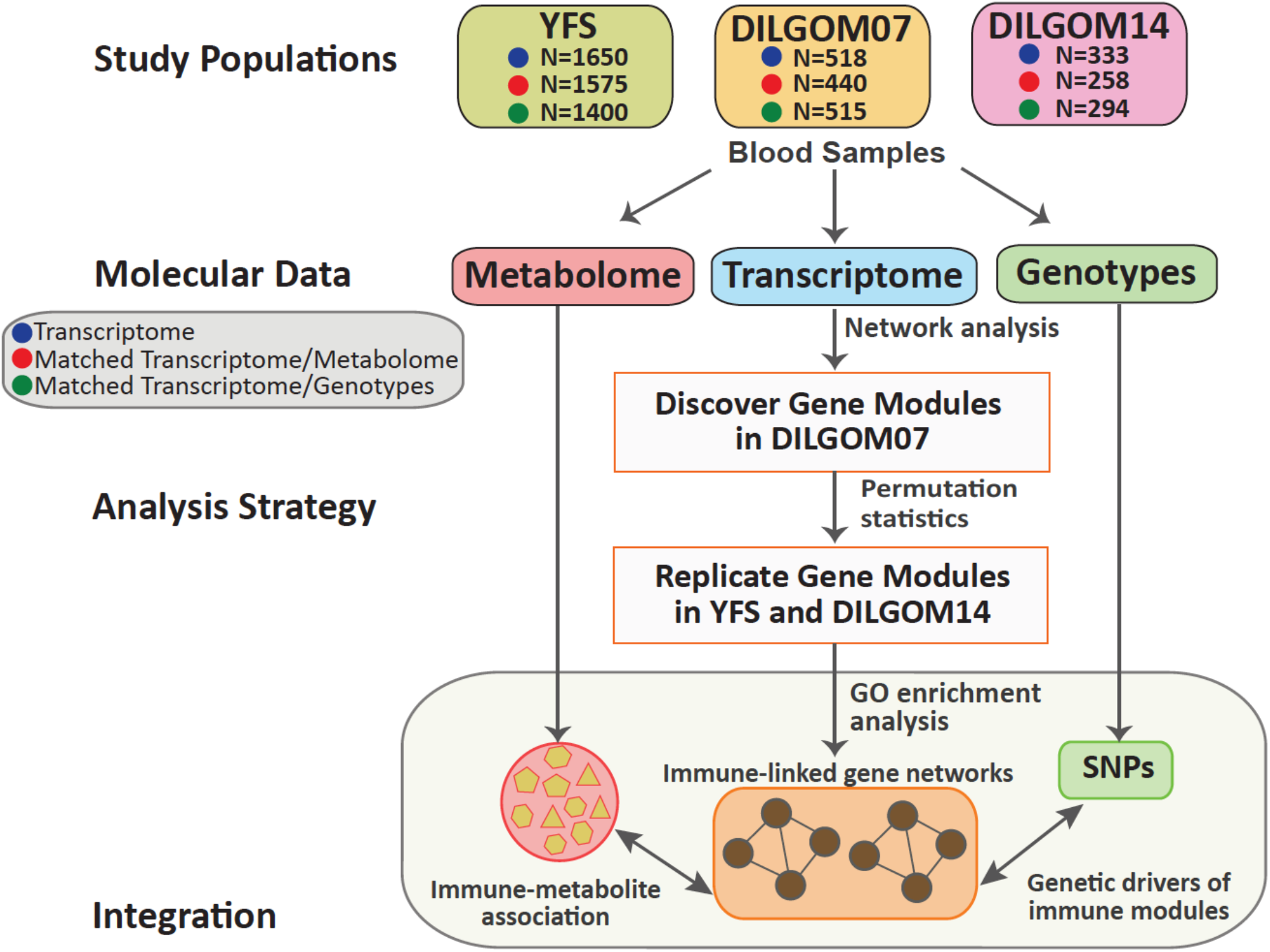
Schematic of the study design.

DILGOM and YFS genotyping was performed using Illumina Human 610 and 670 arrays, respectively, with subsequent genotype imputation performed using IMPUTE2 (17) and the 1000 Genomes Phase I version 3 reference panel. For both cohorts, whole blood transcriptome profiling was performed using Illumina HT-12 arrays and serum metabolomics profiling was carried out using the same serum NMR metabolomics platform (Brainshake, Ltd) (18). Individuals on lipid-lowering medication and pregnant women were excluded from the metabolome analyses (**Methods**). Of the 158 metabolites analysed, 148 were directly quantified and 10 derived (**Table S1**). After filtering, matched transcriptome and metabolome data was available for 440 individuals in DILGOM07 and 216 of these individuals (DILGOM14) who were profiled at seven-year follow-up. In YFS, 1,575 individuals were available with similar data (see **Methods** for details).

### Robust immune gene coexpression networks from blood

We first identified networks of tightly coexpressing genes in DILGOM07 and then used a permutation approach, NetRep (19), to statistically test replication patterns of density and connectivity for these networks in YFS. For module detection, we applied weighted gene coexpression network analysis (WGCNA) to all 35,422 probes in the DILGOM07 data, identifying a total of 40 modules of coexpressed genes (**Methods**). For each module, we used NetRep to calculate seven preservation statistics in the YFS, generate empirical null distributions for each of these test statistics, and calculate their corresponding *P*-values (19,20). A module was considered strongly preserved if the *P* was < 0.001 for all seven preservation statistics (Bonferroni correction for 40 modules). Of the 40 DILGOM07 modules, 20 were strongly preserved in YFS (**Table S2**). For each of the 20 replicated modules, we defined core gene probes, those which are most tightly coexpressed and thus robust to clustering parameters, using a permutation test of module membership (**Methods** and **Table S3**).

To identify modules of putative immune function, we carried out Gene Ontology (GO) biological process enrichment analysis using GOrilla for the core genes of each replicated module (21). Significant GO terms (FDR < 0.05) were then summarised into representative terms based on semantic similarity using REVIGO (22) (**Figure S1**). A module was considered immune related if it was significantly enriched for GO terms “immune system processes” (GO: 0002376) and/or “regulation of immune system processes” (GO:0002682) in the REVIGO output. Six out of 20 modules were enriched for at least one of these terms (**Table S4**). We also identified two additional modules, which were not enriched for any GO terms, but have been previously linked to immune functions related to mast cell and basophil function (13) and platelet aggregation activity (23). The eight modules encoded diverse immune functions, including cytotoxic, viral response, B cell, platelet, neutrophil, mast cell/basophil, and general immune-related functions. Each immune module's gene content and putative biological function is summarized in **Table 1.**

**Table 1:** Immune module gene content and putative biological function based on GO terms (top three shown) and literature.

| Module | Size | GO terms | Literature-based immune related function of genes |
| --- | --- | --- | --- |
| Cytotoxic cell-like module (CCLM) | 130<br>(115) | Immune system process<br>Defense response<br>Immune response | Cytotoxic effectors (*GZMA*, *GZMB*, *GZMM*, *CTSW*, *PRF1* (66)); surface receptors (*IL2RB*, *SLAMF6*, *CD8A*, *CD8B*, *CD2*, *CD247*, *KLRD1*, *KLRG1* (66–68)); T and NK cell differentiation (*ID2* and *EOMES* (69,70)), activation (*ZAP70* and *CBLB* (71,72)), and recruitment (*CX3CR1*, *CCL5*, *CCL4L2* (73)). |
| Viral response module (VRM) | 95<br>(88) | Response to virus<br>Type I interferon signaling pathway<br>Response to biotic stimulus | Type I interferon-induced antiviral activity (*IFITM1*, *IFIT1*, *IFIT2*, *IFIT3*, *IFIT5*, *IFI44*, *IFI44L*, *IFI6*, *MX1*, *ISG15*, *ISG20*, *HERC5* (74,75)); viral RNA degradation (*OAS1*, *OAS2*, *OAS3*, *OASL*, *DDX60* (31)); type 1 interferon-signalling pathway (*IRF9*, *STAT1*, *STAT2* (76,77)). |
| B cell activity module (BCM) | 54<br>(49) | Immune system process<br>Immune response<br>B cell activation | B cell surface markers (*CD79A*, *CD79B*, *CD22* (34,78)); B cell activation (*BANK1*, *BTLA*, *CD40*, *TNFRSF13B*, *TNFRSF13C* (79)), development (*POU2AF1*, *BCL11A*, *RASGRP3* (80)), migration (*CXCR5*, *CCR6* (80,81)), and their regulation (*CD83*, *FCER2*, *FCRL5* (82)); antigen presentation (*HLA-DOA*, *HLA-DOB* (83)). |
| *Platelet module (PM) | 114<br>(106) | Coagulation<br>Blood coagulation<br>Cell activation | Platelet receptor signalling, activation, and coagulation (*GP6*, *GP9*, *ITGA2B*, *ITGB3*, *ITGB5*, *MGLL*, *MPL*, *MMRN1*, *PTK2*, *VCL*, *THBS1*, *F13A1*, *VWF*, (84)); regulating platelet activity (*SEPT5*, *TSPAN9* (85,86)). |
| *Neutrophil module (NM) | 26<br>(26) | Killing of cells of other organism<br>Cell killing<br>Response to fungus | Anti -microbial, -fungal, and -viral activity (*DEFA1*, *DEFA1B*, *DEFA3*, *DEFA4*, *ELANE*, *BPI*, *RNASE2*, *RNASE3* (87–90)); neutrophil mediated activity (*AZU1*, *LCN2*, *MPO*, *CEACAM6*, *CEACAM8*, *OLFM4* (90,91)) and its regulation (*LCN2*, *CAMP*, *OLR1* (50,92,93)) |
| *Lipid-leukocyte module (LLM) | 13<br>(13) | **Mast cell and basophil function | Mast cell and basophil related immune response and allergic inflammation (*FCER1A*, *HDC*, *GATA2*, *SLC45A3*, *CPA3*, *MS4A3* (13,94,95)) |
| General immune module A (GIMA) | 509<br>(482) | Immune system process<br>Defense response<br>Regulation of response to stimulus | These modules contain genes involved in a broad range of immune processes and their regulation such as signalling; cell death; defense response to stress, inflammation, and external stimuli; leukocyte activation, migration, and adhesion. |
| General immune module B (GIMB) | 74<br>(69) | Immune response-activating signal transduction<br>Positive regulation of immune response<br>Activation of immune response |  |

### Immune module association analysis for eQTLs and metabolite levels

For each gene module, we performed a genome-wide scan to identify module QTLs (mQTLs) that regulate expression. In DILGOM07 and YFS, the module eigengene was regressed on each SNP, then mQTL test statistics were combined in meta-analysis (**Methods**). Significant mQTLs were further examined at individual gene expression levels. A genome-wide significance level (*P-*value < 5x10^-8^) was used to identify mQTLs and significant *trans* effects on individual gene expression (**Figure 2** and **Table 2**). Leukocyte and platelet counts were available for YFS and were used to test the robustness of module associations with mQTLs and metabolites. Six modules showed statistically significant association with platelet or leukocyte counts (*P*-value < 0.05) (**Table S5**), however adjustment for leukocyte counts did not affect mQTL nor metabolite-module associations, with the exception of the PM and CCLM discussed below (**Table S6).** Since we did not have cell counts available for DILGOM07, all the immune-metabolite associations discussed below, unless otherwise noted, have not been adjusted for cell counts.

**Figure 2:**
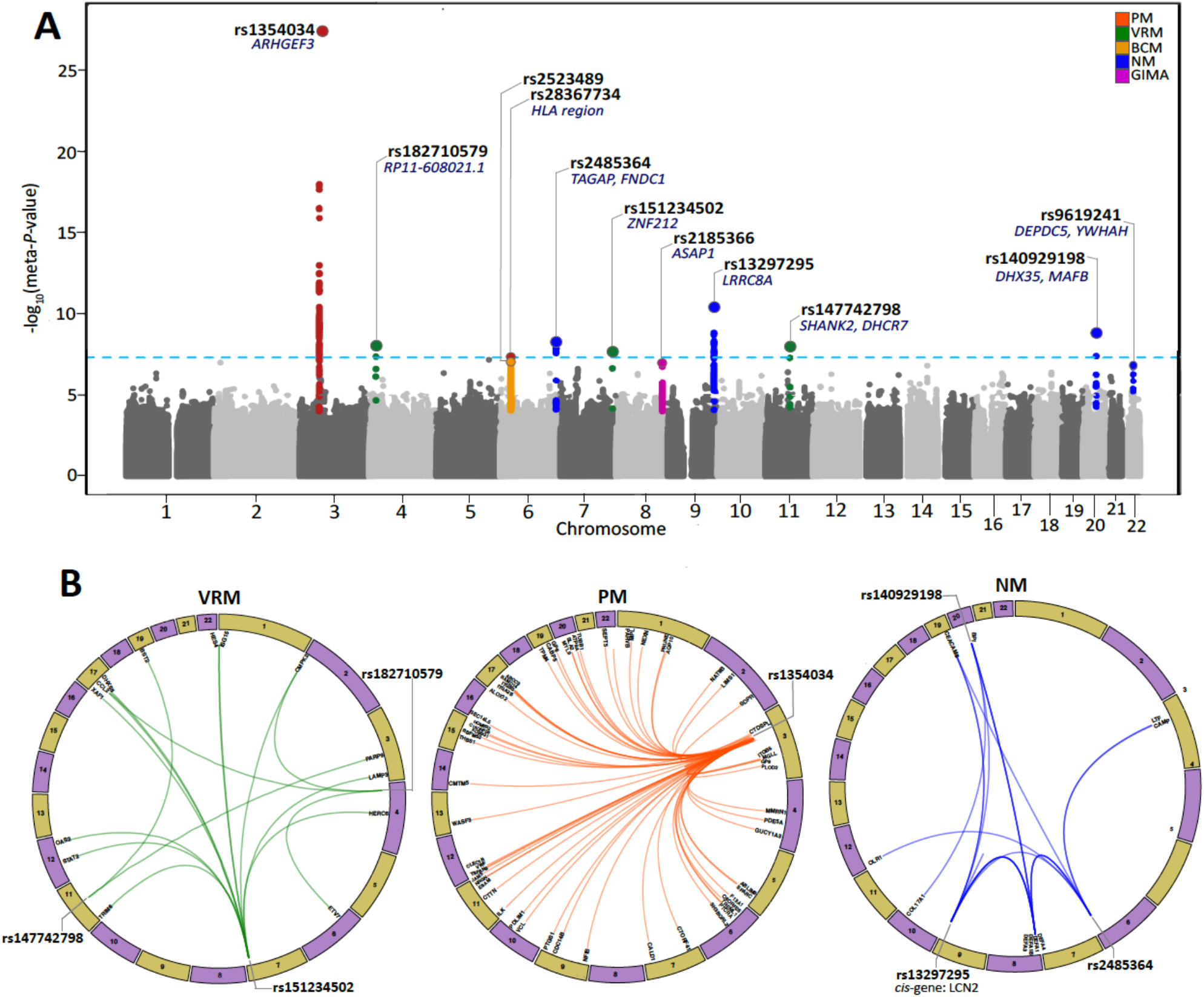
Module and expression QTL analysis. (**A**) Manhattan plot of meta-analysed *P*-values from the DILGOM/YFS module QTL analysis. The lead SNP and its closest genes are noted. Each significant mQTL locus is coloured by module. The horizontal dashed line represents genome-wide (meta *P*-value < 5 x10^-8^) significance. (**B**) Circular plot summarising the individual gene associations (meta-*P*-value < 5 x10^-8^) for the lead module QTLs in the VRM, PM and NM modules. Lead SNPs and *cis* genes are labeled outside the ring. PM (platelet module), VRM (viral response module), CCLM (cytotoxic cell-like module), NM (neutrophil module), BCM (B cell activity module), and GIMA (General immune module A).

**Table 2:** QTLs for immune gene modules. Modules: VRM (viral response module), BCM (B cell activity module), PM (platelet module), NM (neutrophil module), GIMA (General immune module A).

| Module | Top SNP | CHR | Hg19 Pos.<br>(Mb) | Allele<br>(minor/major) | MAF<br>(Avg) | P-value<br>DILGOM07<br>(effect size) | P-value<br>YFS<br>(effect size) | Meta<br>P-value |
| --- | --- | --- | --- | --- | --- | --- | --- | --- |
| VRM | rs182710579 | 4 | 19768086 | G/T | 0.012 | $2.01 \times 10^{-4}$ (0.05) | $8.10 \times 10^{-6}$ (0.02) | $9.23 \times 10^{-9}$ |
| | rs151234502 | 7 | 148950168 | T/C | 0.012 | $2.59 \times 10^{-1}$ (0.01) | $5.31 \times 10^{-9}$ (0.03) | $2.46 \times 10^{-8}$ |
| | rs147742798 | 11 | 70947761 | T/C | 0.016 | $1.51 \times 10^{-3}$ (0.04) | $1.66 \times 10^{-6}$ (0.02) | $9.43 \times 10^{-9}$ |
| BCM | rs2523489 | 6 | 31348878 | T/C | 0.186 | $1.42 \times 10^{-1}$ (0.005) | $5.29 \times 10^{-8}$ (0.006) | $6.27 \times 10^{-8}$ |
| PM | rs1354034 | 3 | 56849749 | T/C | 0.284 | $7.11 \times 10^{-14}$ (-0.02) | $1.51 \times 10^{-16}$ (-0.008) | $7.35 \times 10^{-28}$ |
| | rs28367734 | 6 | 3128657 | A/G | 0.108 | $5.40 \times 10^{-4}$ (0.02) | $2.02 \times 10^{-5}$ (0.006) | $5.44 \times 10^{-8}$ |
| NM | rs2485364 | 6 | 159512260 | C/T | 0.466 | $1.78 \times 10^{-3}$ (0.009) | $6.05 \times 10^{-7}$ (0.004) | $3.93 \times 10^{-9}$ |
| | rs13297295 | 9 | 131659724 | C/T | 0.085 | $4.26 \times 10^{-2}$ (0.009) | $8.39 \times 10^{-11}$ (0.01) | $3.93 \times 10^{-11}$ |
| | rs140929198 | 20 | 38555870 | A/G | 0.031 | $2.98 \times 10^{-2}$ (0.03) | $8.47 \times 10^{-9}$ (0.01) | $1.41 \times 10^{-9}$ |
| | rs2185366 | 22 | 32317905 | G/A | 0.414 | $8.06 \times 10^{-3}$ (-0.007) | $5.16 \times 10^{-6}$ (-0.004) | $1.36 \times 10^{-7}$ |
| GIMA | rs2185366 | 8 | 131342722 | T/C | 0.421 | $2.0 \times 10^{-2}$ (0.007) | $1.52 \times 10^{-6}$ (0.004) | $1.05 \times 10^{-7}$ |

### Cytotoxic cell-like module (CCLM)

CCLM was associated with 24 metabolites, mainly consisting of fatty acids, intermediate density lipoproteins, and CRP (**Figure 3** and **Table S7**). The top associated metabolite was the average number of double bonds in a fatty acid chain (meta-*P-*value = 7.23 x 10^-7^). It is known that CRP augments cytotoxic responses by binding to NK cells, modulating their activity (24), enhancing cytotoxic responses of NK cells against tumour cells (25), and sensitizing endothelial cells to cytotoxic T-cell mediated destruction (26). The interaction between fatty acid double bond saturation and anti-inflammatory response is well characterised, and unsaturated fatty acids have been shown to induce cytotoxicity in *in vitro* cancer cell lines as well as animal models of tumour incidence and growth (27,28). Adjustment of CCLM-metabolite associations for leukocyte counts resulted in the gain of 38 additional associations and loss of four (creatinine, ratio of polyunsaturated fatty acids to total fatty acids, VLDL particle size, and CRP) existing associations (**Table S6**). CCLM had no significant mQTLs.

**Figure 3:**
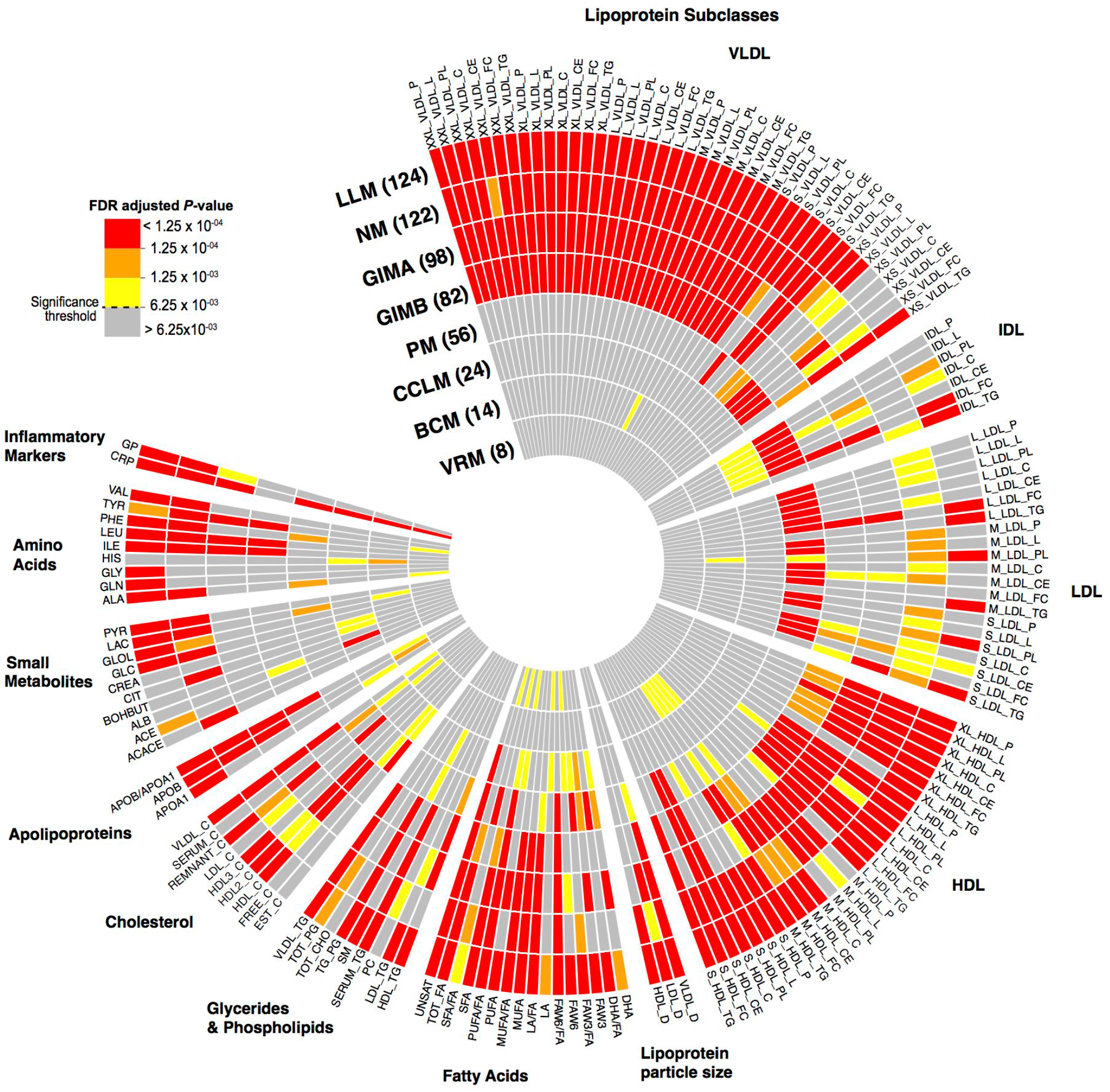
Metabolite associations with immune gene modules. Circular heatmap of associations between individual metabolites and the module eigengene of each module (coloured by FDR-adjusted *P-*values). Concentric circles represent modules, with numbers in parentheses denoting total number of metabolites associated with that module at FDR-adjusted *P*-value < 6.25 x 10^-3^. NM (neutrophil module), LLM (lipid leukocyte module), GIMA, and GIMB (General immune modules A and B), PM (platelet module), CCLM (cytotoxic cell-like module), BCM (B cell activity module), and VRM (viral response module). See **Table S1** for full metabolite descriptions.

### Viral response module (VRM)

Three genome-wide significant mQTLs were identified for the VRM (**Figure 2** and **Table 2**). The strongest mQTL, rs182710579 (meta*-P*-value = 9.22 x 10^-9^), is within a known lincRNA locus (RP11-608O21.1) (**Figure S2A**). Rs182710579 was a *trans* eQTL for 3 genes in the VRM (**Table S8**). The strongest association was seen with *CCL2* (meta-*P*-value = 6.78 x 10^-12^), a pro-inflammatory chemokine involved in leukocyte recruitment during viral infection (29,30). The next strongest mQTL, rs151234502, resides within intron 4 of the relatively unstudied *ZNF212*, part of a zinc finger gene cluster at 7q36 (**Figure S2B**). Rs151234502 modulated expression of 11 VRM genes *in trans* (**Table S8**). The strongest association was with *OAS2* (meta-*P*-value = 8.98 x 10^-10^), an interferon-induced gene encoding an enzyme promoting RNase L-mediated cleavage of viral and cellular RNA (31). The third mQTL, rs147742798, was an intergenic SNP located between *SHANK2* and *DHCR7* at 11q13.4 (**Figure S2C**). Rs147742798 was a *trans* eQTL for 2 genes in the VRM, *BST2* and *PARP9* (**Table S8**). *BST2* encodes a trans membrane protein with interferon-inducible antiviral function (32). Studies have previously shown induction of fatty acid biosynthesis by a range of viruses (33). VRM was associated with eight metabolites, including amino acids (alanine, phenylalanine), fatty acids (omega-6 fatty acids, polyunsaturated fatty acids, saturated fatty acids, and total fatty acids), and cholesterol esters in medium VLDL (**Figure 4** and **Table S7**). Consistent with its putative role in viral response, VRM was strongly associated with CRP (meta *P*-value = 2.38 x 10^-10^).

**Figure 4:**
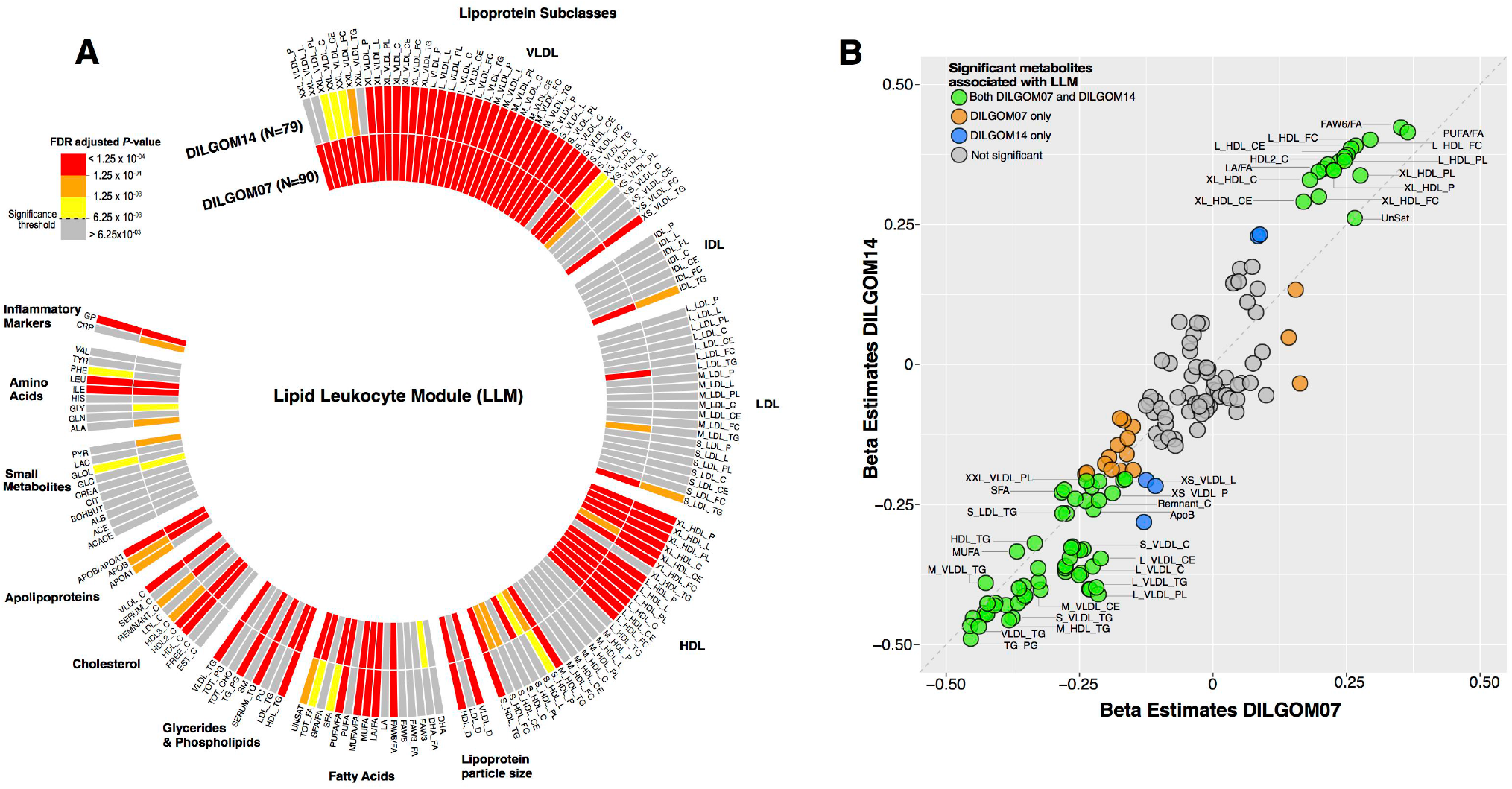
Temporally stable metabolite associations with the LLM. (**A**) Circular heatmap for association between individual metabolites and the LLM. (**B**) Comparison of the effect size estimates of metabolite association with LLM in DILGOM07 and DILGOM14 shows that the overall association patterns are consistent across the two time-points. Colours denote metabolites that are significantly associated with the LLM in DILGOM07 only (orange), DILGOM14 only (blue), and across both time-points (green). The grey dashed line is the x=y line.

### B cell activity module (BCM)

The BCM was associated with 14 metabolites including CRP, histidine, lactate, apolipoprotiens, and mainly medium HDL subclass of lipoproteins. (**Figure 3** and **Table S7**). The strongest association was seen with CRP (meta-*P*-value = 2.65 x 10^-8^). Histidine, the second most strongly associated metabolite, is catabolized to histamine by histidine decarboxylase. The relationship between B cells and histamine is a central part of the allergic reaction where IgE released by B cells blankets mast cells, causing them to release histamine. While no mQTLs for BCM reached genome-wide significance, there was some evidence in the YFS for the MHC class I locus (**Figure 2** and **Table 2**). The top signal was located between *HLA-B/C* and *MICA* (rs2523489, meta-*P*-value = 6.27 x 10^-8^) (**Figure S3**). The HLA class I region is well-known to be associated with autoimmune diseases, where the role of B cells is well recognized Rs2523489 was a *trans* eQTL for *CD79B* (meta-*P*-value = 1.16 x 10^-9^), a subunit of the antigen-binding B cell receptor complex (34).

### Platelet module (PM)

PM had the strongest mQTL of any gene module, an intronic SNP of the *ARHGEF3* gene at 3p14.3 (rs1354034; meta-*P*-value = 7.35 x 10^-28^, **Figures 2** and **S4A, Table 2**). *ARHGEF3* encodes a Rho guanine nucleotide exchange factor, a catalyst of Rho GTPase conversion from inactive GDP-bound to active GTP–bound form. Rs1354034 was an eQTL for the majority of genes in the PM, all of which were *in trans*. An intergenic SNP, rs2836773 (meta-*P*-value = 5.4 x 10^-8^), at the HLA locus was also identified as an mQTL for PM (**Figure S4B**). The *ARHGEF*3 mQTL (rs1354034) exhibited a strong *trans*-regulatory effect and was associated with 61 PM genes (65 unique probes) (**Table S8**). The top *trans* eQTL was *ITGB3* (meta-*P*-value = 5.09 x 10^-42^), a gene encoding the β_3_ subunit of the heterodimeric integrin receptor (integrin α_IIb_β_3_). This integrin receptor is most highly expressed on activated platelets and plays a key role in mediating platelet adhesion and aggregation upon binding to fibrinogen and Willebrand factor (35,36). Our data are consistent with previous observations of the diverse *trans* eQTL effects of rs1354034 (23), including the putative splice-QTL effects of rs1354034 on *TPM4*, a significant eGene in the PM.

*ARHGEF3* itself is of intense interest to platelet biology. It has previously been shown that silencing of *ARHGEF3* in zebrafish prevents thrombocyte formation (37). To test whether *ARHGEF3* expression had an effect on PM genes, we regressed out *ARHGEF3* levels and re-ran the eQTL analysis. Adjusting for *ARHGEF3* did not attenuate the *trans*-associations of rs1354034, suggesting either independence of downstream function for *ARHGEF3* and rs1354034 or post-transcriptional modification of ARHGEF3. Previous GWAS studies have shown rs1354034 is associated with platelet count and mean platelet volume (37), however, perhaps due to power, we found no significant relationship between platelet counts and rs1354034 in YFS. While platelet counts were positively associated with the PM (β = 0.29; *P*-value = 8.23 x 10^-30^) (**Table S5**), the association between rs1354034 and the PM was still highly significant when conditioning on platelet counts (β = -0.33; *P*-value = 1.40 x 10^-17^).

PM displayed diverse metabolic interactions and was associated with 55 metabolites, largely comprising of lipoprotein subclasses and fatty acids, as well as CRP (**Figure 3** and **Table S7**). Cholesterol esters in small HDL particles were most strongly associated with the PM (meta-*P*-value = 9.45 x 10^-20^). HDL has been shown to exhibit antithrombotic properties by modulating platelet activation, aggregation and coagulation pathway (38). On the other hand, pro-atherogenic lipoproteins effects on platelets has been recognised as an important driver in development of atherosclerosis. For example, LDL has been shown to influence platelet activity either by enhancing platelet responsiveness to aggregating stimuli or inducing aggregation (39,40). Moreover, LDL specific binding sites on platelets have also been reported (41,42). As noted above, the PM was associated with platelet counts, and adjustment of PM-metabolite associations for platelet counts in the YFS resulted in attenuation of approximately half of the weakest metabolite associations, however the strongest were maintained (**Table S6**). Association with VLDL particle size and three others were gained following the adjustment (**Table S6**).

### Neutrophil module (NM)

Three loci were identified as mQTLs for NM (**Figure 2** and **Table 2**). The top mQTL was intronic to *LRRC8A* at 9q34.11 (rs13297295; meta-*P*-value = 3.93 x 10^-11^, **Figure S5A**). *LRRC8A* encodes for a trans-membrane protein shown to play a role in B and T cell development and T cell function (43,44). Two additional intergenic mQTLs were located at the *TAGAP* locus at 6q25.3 (rs2485364; meta-*P*-value = 3.93 x 10^-9^) and at 20q12 (rs140929198; meta-*P*-value = 1.41 x 10^-9^) (**Figures S5B-C**). Rs13297295 was a strong *trans* regulator of NM and was an eQTL for 8 NM genes (10 unique probes), in particular the major alpha defensins (*DEFA1-DEFA4*), the genes of highest centrality in the module (**Table S8**). Rs13297295 was a *cis*-eQTL for another core NM gene, *LCN2* (permuted meta-*P*-value = 1 x 10^-4^) (**Table S8**). *LCN2* is expressed in neutrophils and inducible by TLR activation, acting as an antimicrobial agent via sequestration of bacterial siderophores to prevent iron uptake (45–47). *LCN2*'s role in acute phase response appears to be related to cardiovascular diseases, such as heart failure (48). At the *TAGAP* locus, rs2485364 was a *trans*-eQTL for 8 NM genes (10 probes) and was also a strong driver of *LCN2* (meta-*P*-value = 9.11 x 10^-17^) (**Table S8**). Consistent with our findings, neutrophils from *LCN2* deficient mice have been shown to have impaired chemotaxis, phagocytic capability, and increased susceptibility to bacterial and yeast infections compared to wild type (49,50). This suggests a possible functional role of *TAGAP* variants in regulating neutrophil migration through *LCN2*.

NM was associated with 121 circulating metabolites (~76% of all metabolites analysed) as well as CRP (**Figure 3** and **Table S7**). The strongest is the previously reported association with inflammatory biomarker GlycA (meta-*P*-value = 2.68 x 10^-25^) (10), however NM's association with various lipoprotein subclasses, particle sizes of lipoproteins, fatty acids, cholesterol, apolipoproteins, glycerides and phospholipids, amino acids, and other small molecules indicates it has a potentially major role in linking neutrophil function to metabolism.

### Lipid-Leukocyte module (LLM)

Together with NM, the LLM showed extensive metabolic associations. Overall, 123 metabolites and CRP were associated with LLM, with the strongest being the ratio of triglycerides to phosphoglycerides (meta-*P*-value = 5.16 x 10^-138^, **Figure 3**, **Table S7**). With the inclusion of the YFS, these findings strongly replicate previous LLM-metabolite associations (14) as well as highlight additional metabolite associations. We also confirm the previous strong negative association between CRP and LLM (meta-*P*-value = 8.16 x 10^-20^). Consistent with previous studies, no mQTLs were detected for LLM.

### General immune modules A and B (GIMA and GIMB)

No mQTLs were associated with GIMA and GIMB, however these modules were associated with 97 and 82 metabolites, respectively (**Figure 3** and **Table S7**). Cholesterol esters in small HDL and the mean diameter for VLDL particles exhibited the strongest associations with GIMA (meta-*P*-value = 1.56 x 10^-30^) and GIMB (meta-*P*-value = 1.83 x 10^-15^), respectively. The GIMA was also associated with CRP (meta-*P*-value = 5.7 x 10^-5^). Other metabolite associations with these two modules include mainly the VLDL and HDL subclass of lipoproteins and fatty acids, however, due to their large size and heterogeneous composition, interpretation of metabolic relationships of GIMA and GIMB is limited.

### Long-term stability of interactions between metabolites, immune gene modules and mQTLs

The 216 individuals in both the DILGOM 2007 and 2014 follow-up allowed investigation of the long-term stability of immunometabolic and mQTL relationships. Across this seven-year period, the eight immune gene coexpression networks were strongly preserved (all preservation statistics' permutation *P*-values <0.001; **Table S9**). The metabolite-metabolite correlation structure was also largely consistent between DIGOM07 and DILGOM14 (**Figure S6**).

Next, we examined how metabolite interactions with immune gene modules changed over the 7-year time period (**Methods**). The LLM-metabolite associations were the most consistent over time with 90 and 79 metabolites reaching significance in DILGOM07 and DILGOM14, respectively, of which 74 were significant at both time points (**Figure 4A**, **Table S10**). The direction and effect size of LLM-metabolite associations were largely maintained (**Figure 4B**). For the neutrophil module, only the pyruvate association was significantly maintained over time, however there was some evidence that other expected NM-metabolite associations were stable over time, including GlycA (**Table S10**). While no metabolite associations were significantly maintained for platelet module, rs1354034 was a temporally stable mQTL of PM (mQTL *P*-value = 4.87 x 10^-7^). No other mQTLs reached significance for temporal stability.

## Conclusions

This study has utilised over 2,000 individuals to map the immuno-metabolic crosstalk operating in circulation. We have identified and characterised eight robust immune gene modules, their genetic control and interactions with diverse metabolites, including many of clinical significance (e.g. triglycerides, HDL, LDL, branched-chain amino acids). Furthermore, our findings are consistent with and build upon those of previous studies. In addition to five newly identified gene modules, their mQTLs and metabolite interactions, we have replicated the previously characterised LL module and confirm its association with lipoprotein subclasses, lipids, fatty acids, and amino acids (13,14). Associations between the core genes in the LL module and isoleucine, leucine, and various lipids were also identified independently in the KORA cohort (12). Importantly, we have shown the long-term stability of LL and neutrophil module coexpression and metabolite interactions, and we have greatly expanded the number of known biomarkers associated with the NM from one (GlycA) to 123 (10). Our study has also expanded the widespread *trans* eQTL effects at the *ARHGEF3* locus (23), shows them to be strongly maintained within individuals over time, and further identifies extensive lipoprotein and fatty acid metabolite interactions that may be a consequence of these *trans* effects.

Taken together, our analyses illustrate the rapidly growing body of evidence intimately linking the immunoinflammatory response to the blood metabolome. With finer-resolution maps of these interactions, new biomarkers of chronic and acute inflammatory states are likely to emerge. With *in vivo* and interventional studies, modulation of these metabolite-immune interactions through existing lipid-lowering medications, gut microbe effects or dietary changes may provide new ways the immune system itself can be utilised to lessen the burden of cardiometabolic disease.

## Methods

### Study Populations

This study used data from two population-based cohorts, the Dietary, Lifestyle, and Genetic determinant of Obesity and Metabolic syndrome (DILGOM; N=518) and the Cardiovascular Risk in Young Finns Study, (YFS; N=1,650), which have been described in detail elsewhere (13,51). All subjects enrolled in these studies gave written informed consent.

The DILGOM study is a subsample of the FINRISK 2007 cross-sectional population-based survey, which recruited a random sample of 10,000 individuals between 25–74 years of age, stratified by sex and 10-year age groups, from five study areas in Finland. All 6258 individuals who participated in the FINRISK 2007 baseline health examination were invited to attend the DILGOM study (N=5,024), 630 of whom underwent at least one of the genotyping, transcriptomics or metabolomics profiling considered here. Ethics approval was given by the Coordinating Ethical Committee of the Helsinki and Uusimaa Hospital District. In 2014, a follow-up study was conducted, for which 3,735 individuals from the original study re-participated. Samples collected in 2007 and 2014 are referred to as DILGOM07 and DILGOM14, respectively.

The DILGOM study is a cross-sectional population-based survey conducted in 2007, which randomly recruited 5,325 unrelated individuals aged between 25–74 years of age from the Helsinki region of Finland, 630 of whom underwent at least one of the genotyping, transcriptomics or metabolomics profiling considered here. Ethics approval was given by the Coordinating Ethical Committee of the Helsinki and Uusimaa Hospital District. In 2014, a follow-up study was conducted, for which 1,273 individuals from the original study re-participated. Samples collected in 2007 and 2014 are referred to as DILGOM07 and DILGOM14, respectively.

The YFS is a longitudinal prospective cohort study that started in 1980, with follow-up studies carried out every three years, to monitor cardiovascular disease risk factors in children and adolescents from five major regions of Finland (Helsinki, Kuopio, Turku, Oulu, and Tampere). In the baseline study a total of 3,596 children and adolescents in age groups 3, 6, 9, 12, 15, and 18 years participated, who were randomly selected from the national public register, details of which are described in (51). In this current study, data collected from the 2011 follow-up study (participants aged 34, 37, 40, 43, 46, and 49 years) were analysed. Ethics approval for the study research protocols was given by the Joint Commission on Ethics of Turku University and Turku University Central Hospital.

### Sample Collection

Venous blood was collected following an overnight fast in all three studies. Samples were centrifuged, the resulting plasma and serum samples were aliquoted into separate tubes and stored at −70°C for analyses. Protocols for the blood sampling, physiological measurements, and clinical survey questions were similar across the YFS and DILGOM studies, and are described extensively in (13,52).

### Genotyping and Imputation

Whole blood genomic DNA obtained from both cohorts was genotyped using the Illumina 610-Quad SNP array for DILGOM07 (N=555) (13) and a custom generated 670K Illumina BeadChip array for YFS (N=2,443) (53). The 670K array shares 562,643 SNPs with the 610-quad array. The 670K array removes poorly performing SNPs from the 610-quad array and improves copy number variation coverage (53). Genotype calling was performed with the Illuminus clustering algorithm (54). Quality control was as previously described in (13) and (53) for DILGOM and YFS, respectively. Genotypes were imputed to the 1000 Genomes Phase 1 version 3 reference panel using IMPUTE2 in both DILGOM and YFS (17). Poorly imputed SNPs based on low call-rate (< 0.90 for DILGOM, < 0.95 for YFS), low-information score (< 0.4), minor allele frequency < 1%, and deviation from Hardy-Weinberg equilibrium (*P* < 5 x 10^-6^) were then removed. A total of 7,263,701 SNPs in DILGOM and 6,721,082 in YFS passed quality control, with 6,485,973 common between the two. A total of N=518 samples in DILGOM and N=2,443 samples in YFS individuals passed quality control filters.

### Metabolomics profiling

Metabolite concentrations for DILGOM07 (N=4,816), DILGOM14 (N=1,273), and YFS (N=2,046) were quantified from serum samples utilizing a high-throughput NMR metabolomics platform (Brainshake Ltd, Helsinki, Finland) (18,55). Details of the experimental protocol including sample preparation, NMR spectroscopy and metabolite identification has been previously described in (13,18). A total of 158 metabolite measures were assessed, of which 148 were directly measured and 10 were derived (**Table S1**). The 148 measures include the constituents of 14 lipoprotein subclasses (98 measurements total), sizes of 3 lipoprotein particles, 2 apolipoproteins, 8 fatty acids, 8 glycerides and phospholipids, 9 cholesterols, 9 amino acids, 1 inflammatory marker, and 10 small molecules (involved in glycolysis, citric acid cycle and urea cycle). The lipoprotein subclasses are classified according to size as follows: chylomicrons and extremely large VLDL particles (average particle diameter at least 75 nm); five VLDL subclasses: very large VLDL (average particle diameter of 64.0 nm), large VLDL (53.6 nm), medium VLDL (44.5 nm), small VLDL (36.8 nm), and very small VLDL (31.3 nm); intermediate-density lipoprotein (IDL) (28.6 nm); three LDL subclasses: large LDL (25.5 nm), medium LDL (23.0 nm), and small LDL (18.7 nm); and four HDL subclasses: very large HDL (14.3 nm), large HDL (12.1 nm), medium HDL (10.9 nm), and small HDL (8.7 nm). Measurements with very low concentration, set as zero by the NMR pipeline, were set to the minimum value of that particular metabolite. Measurements rejected by automatic quality control or with detected irregularities were treated as missing. Undefined derived ratios arising from measurements with very low concentration (i.e. zero) were also treated as missing. Measurements were log2 transformed to approximate a normal distribution.

C-reactive protein (CRP), an inflammatory marker, was quantified from serum using a high sensitivity latex turbidimetric immunoassay kit (CRP-UL assay, Wako Chemicals, Neuss, Germany) and an automated analyser (Olympus AU400) in DILGOM07 (N=5000), DILGOM14 (N=1308), and YFS (N=2046). CRP levels were log2 transformed.

### Gene expression, processing and normalization

Transcriptome-wide gene expression levels were quantified by microarrays from peripheral whole blood using similar protocols in all three cohorts, and have been previously described for DILGOM07 (13) and YFS (56). Stabilised total RNA was obtained from whole blood using a PAXgene Blood RNA System and the protocols recommended by the manufacturer. In DILGOM07, RNA integrity and quantity was evaluated using an Agilent 2100 Bioanalyzer. In YFS, RNA integrity and quantity were evaluated spectrophotometrically using an Eppendorf BioPhotomer and the RNA isolation process was validated using an Agilent RNA 6000 Nano Chip Kit. RNA was hybridized to Illumina HT-12 version 3 BeadChip arrays in DILGOM07 and to Illumina HT-12 version 4 BeadChip arrays in DILGOM14 and YFS.

For DILGOM07, data was preprocessed as described in Inouye *et al.* (13). Briefly, for each array the background corrected probes were subjected to quantile normalization at the strip-level. Technical replicates were combined by bead count weighted average and replicates with Pearson correlation coefficient < 0.94 or Spearman’s rank correlation coefficient < 0.60 were removed. Expression values for each probe were then log2 transformed. For YFS, background corrected probes were subjected to quantile normalization followed by log2 transformation. For DILGOM14, probes matching to the erythrocyte globin components (N=4) and those that hybridized to multiple locations spanning more than 10Kb (N=507) were removed. Probes with average bead intensity of 0 were treated as missing. The average bead intensity was then log2 transformed and quantile normalized. A total of 35,425 (for DILGOM07), 36,640 (for DILGOM14) and 37,115 (for YFS) probes passed quality control.

### Gene co-expression network analysis and replication

Gene co-expression network modules were identified in DILGOM07 (N=518 individuals with gene expression data) as previously described (10) using WGCNA version 1.47 (57,58) on all probes passing quality control. Briefly, probe co-expression was calculated as the Spearman correlation coefficient between each pair of probes, adjusted for age and sex. The weighted interaction network was calculated as the element-wise absolute co-expression exponentiated to the power 5. This power was selected through the scale-free topology criterion (57), which acts as a penalization procedure to enhance differentiation of signal from noise. Probes were subsequently clustered hierarchically (average linkage method) by topological overlap dissimilarity (57) and modules were detected through dynamic tree cut of the resulting dendrogram with default parameters and a minimum module size of 10 probes (59). Similar modules were merged together in an iterative process in which modules whose eigengenes clustered together below a height of 0.2 were joined. Module eigengenes, representative summary expression profiles, were calculated as the first eigenvector from a principal components analysis of each module’s expression data.

Module reproducibility and longitudinal stability were assessed in YFS (N=1,650 with gene expression data) and DILGOM14 (N=333 with gene expression data) respectively using the NetRep R package version 0.30.1 (19). Briefly, a permutation test (20,000 permutations) of seven module preservation statistics was performed for each module in YFS and DILGOM14 separately. These statistics test the distinguishability and similarity of network features (density and connectivity) for each module in a second dataset (20). Modules were considered reproducible where permutation *P*-values for all seven statistics were < 0.001 (Bonferroni correcting for 40 modules) in YFS, and modules were considered longitudinally stable where *P*-values were < 0.001 for all seven statistics in DILGOM14. Probe co-expression in YFS was calculated as the Spearman correlation coefficient between age and sex adjusted expression levels and the weighted interaction network was calculated as the element-wise absolute co-expression exponentiated to the power 4 as previously described (10). Probe co-expression in DILGOM14 was calculated as the Spearman correlation coefficient between each pair of probes, and the weighted interaction network defined as the element-wise absolute co-expression exponentiated to the power 5.

To filter out genes spuriously clustered into each module by WGCNA we performed a two-sided permutation test on module membership (Pearson correlation between probe expression and the module eigengene) for each reproducible module in DILGOM07 and YFS. Here, the null hypothesis was, for each module, that its probes did not truly coexpress with the module. The null distribution of module membership for each module was empirically generated by calculating the membership between all non-module genes and the module’s eigengene. *P*-values for each probe were then calculated using the following permutation test *P*-value estimator (60):

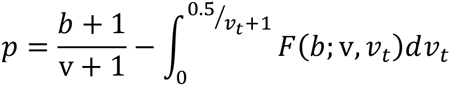

Where *b* is taken as the number of non-module genes with a membership smaller or greater than the test gene’s module membership, whichever number is smaller. *v*, the number of permutations calculated, and *v_t_*, the total number of possible permutations, are both the number of non-module genes. The resulting *P*-value was multiplied by 2 because the test was two-sided. To adjust for multiple testing, false discovery rate (FDR) correction was applied to the *P*-values separately for each module using the Benjamini and Hochberg method (61). We rejected the null hypothesis at FDR adjusted *P*-value < 0.05 in both DILGOM07 and YFS, deriving a subset of core probes for each module.

### Functional annotation of immune modules

Immune modules were identified through over-representation analysis of Gene Ontology (GO) terms in the core gene set for each of the 20 reproducible modules using the web based tool GOrilla (21) with default parameters (performed March 2016). GOrilla was run on two unranked gene lists where core module genes were given as the target list and the background list was given as the 25,233 human RefSeq genes corresponding to any probe(s) passing quality control in both DILGOM07 and YFS. A hypergeometric test was calculated to test whether each module was significantly enriched for genes annotated for each GO term in the “Biological Process” ontology. A GO term was considered significantly over-represented in a module where its FDR corrected *P*-value was < 0.05. FDR correction was applied in each module separately. Significant GO terms for each module were further summarised into a subset of representative GO terms with REVIGO (22) using the RELSIM semantic similarity measure and a similarity cut-off value *C* = 0.5 on genes from *Homo sapiens*. A module was considered to be immune-linked where the representative GO term list contained the parent GO term GO:0002376 (immune system process) and/or GO:0002682 (regulation of immune system processes).

### Statistical Analyses

Reproducible module–metabolite associations were identified through linear regression of each immune module eigengene on each of the 159 metabolites in both DILGOM07 and YFS. Prior to analysis, metabolite data was first subsetted to individuals with matching gene expression profiles, followed by removal of subjects on cholesterol lowering drugs, for YFS (N=62) and DILGOM07 (N=74). Pregnant women in YFS (N=10) and DILGOM (N=2) were further removed from the analysis. A total of 440 individuals in DILGOM07 and 1,575 individuals in YFS had matched gene expression and metabolite data, excluding pregnant women and those individuals taking lipid-lowering medication. Models were adjusted for age, sex, and use of combined oral contraceptive pills. Module eigengenes and metabolite levels were scaled to standard deviation units. To maximize statistical power, a meta-analysis was performed on the DILGOM07 and YFS associations using the fixed-effects inverse variance method implemented in the “meta” R package (https://cran.r-project.org/web/packages/meta/index.html). The meta-*P*-values for the 159 metabolite associations within each module were FDR corrected. An association was considered significant at FDR adjusted *P*-value < 6.25 x 10^-3^ (0.05/8 modules). This Bonferroni adjusted threshold was chosen to further adjust for the multiple modules being tested. To assess the potential confounding effects of blood cell type abundance on metabolite-module association, the model was rerun in YFS adjusting for leukocyte (for CCLM, VRM, BCM, NM, LLM, GIMA, GIMB) and platelet (for PM) counts available for this cohort. The beta values and *P*-values generated with and without adjusting for cell count were then compared. Additionally, to assess the possible effect of cell counts on expression profiles, cell counts were associated with module eigengenes.

Module–metabolite associations were tested for longitudinal stability in DILGOM14 using a linear regression model of each immune module eigengene on each of the 159 metabolites. A total of, 216 individuals in DILGOM had matched gene expression and metabolite data in both 2007 and 2014, after removing pregnant women and individuals on lipid lowering medication at either time point (N=70). Models were adjusted for age and sex. Information on use of oral contraceptives was not available for this cohort. It is worth noting that > 60% of women were more than 50 years old, hence we would expect that rates of contraceptive use would be low and therefore not be a significant confounder. Module eigengenes and metabolite levels were scaled to standard deviation units. An association was considered longitudinally stable where the association was significant (FDR adjusted *P*-value < 6.25 x 10^-3^) in both DILGOM14 and DILGOM07. For sensitivity analysis, the model in DILGOM07 was run without adjusting for oral contraceptive use and this did not affect the significant immune-metabolite associations maintained over the two time-points.

Module quantitative trait loci (mQTLs) were identified through genome-wide association scans with each immune module eigengene using the PLINK2 version 1.90 software (<u>https://www.cog-genomics.org/plink2</u>) (62) in DILGOM07 and YFS. A total of 518 individuals had matched gene expression and genotype data in DILGOM07 and 1400 individuals had matched gene expression and genotype data in YFS. Associations were tested using a linear regression model of each eigengene on the minor allele dosage (additive model) of each SNP. Models were adjusted for age, sex, and the first 10 genetic principal components (PCs). Genetic PCs were generated from a linkage-disequilibrium (LD) pruned set of approximately 200,000 SNPs using flashpca (63). *P*-values for each association in DILGOM07 and YFS were combined in a meta-analysis using the METAL software (64), which implements a sample size weighted Z-score method. A SNP was considered an mQTL if meta-analysis *P*-value (meta-*P*-value) was < 5 x 10^-8^. Blood cell count data available for YFS was utilized to test the robustness of module associations with mQTLs, where the same model was run with and without adjusting for leukocyte and platelet cell counts.

Significant mQTLs were subsequently tested as expression quantitative trait loci (eQTLs) for genes within their respective modules using Matrix eQTL in both DILGOM07 and YFS (65). Both *cis* (mQTL within 1Mb of a given probe) and *trans* (mQTL greater than 5Mb from a given probe or on a different chromosome) associations were tested. Associations were tested using a linear regression model of probe expression on minor allele dosage (additive model) of the mQTL. Models were adjusted for age, sex, and the first 10 genetic PCs. For *trans*-eQTL associations *P*-values in DILGOM07 and YFS were combined in a meta-analysis using the weighted Z-score method and considered significant where the meta-*P*-value < 5x10^-8^. For *cis*-eQTL associations, permutation tests were performed in which gene expression sample labels were shuffled 10,000 times to compute an empirical *P*-value. The permuted model *P*-values and nominal *P*-value were combined across DILGOM and YFS07 in meta-analyses using the weighted Z-score method when computing the permutation test *P*-value. An mQTL was considered a *cis*-eQTL where the permutation test *P*-value < 0.05.

## Acknowledgements

This study was supported by funding from National Health and Medical Research Council of Australia (NHMRC) grant APP1062227. MI was supported by an NHMRC and Australian Heart Foundation Career Development Fellowship (no. 1061435). GA was supported by an NHMRC Early Career Fellowship (no. 1090462). APN and SR were supported by an Australian Postgraduate Award. VS was supported by the Finnish Foundation for Cardiovascular Research. This study was further supported by the Strategic Research Funding from the University of Oulu, Finland, the Sigrid Juselius Foundation, the Academy of Finland (grant numbers 141136, 269517, 283045, 294834, and 297338), the Yrjö Jahnsson Foundation, the Emil Aaltonen Foundation, the Novo Nordisk Foundation, and the Finnish Diabetes Research Foundation. ER was supported by the Academy of Finland (no. 285902). The Young Finns Study has been financially supported by the Academy of Finland: grants 286284 (T.L), 134309 (Eye), 126925, 121584, 124282, 129378 (Salve), 117787 (Gendi), and 41071 (Skidi); the Social Insurance Institution of Finland; Competitive State Research Financing of the Expert Responsibility area of Tampere, Turku and Kuopio University Hospitals (grant X51001); Juho Vainio Foundation; Paavo Nurmi Foundation; Finnish Foundation for Cardiovascular Research; Finnish Cultural Foundation; Tampere Tuberculosis Foundation; Emil Aaltonen Foundation; Yrjö Jahnsson Foundation; Signe and Ane Gyllenberg Foundation; and Diabetes Research Foundation of Finnish Diabetes Association. MJ was supported by the Paulo Foundation, Maud Kuistila Foundation, and Finnish Medical Foundation. The DILGOM study and the National FINRISK study are supported by the Academy of Finland (grant numbers 139 635). The quantitative serum NMR metabolomics platform and its development has been supported by the Academy of Finland, TEKES - the Finnish Funding Agency for Technology and Innovation, the Sigrid Juselius Foundation, and the strategic and infrastructural research funding from the University of Oulu, Finland, as well as by the British Heart Foundation, the Wellcome Trust, and the Medical Research Council, UK. MP is also supported by EU FP7 under grant agreements nr. 313010 (BBMRI-LPC), nr. 305280 (MIMOmics), and HZ2020 633589 (Ageing with Elegans). MAK works in a Unit that is supported by the University of Bristol and UK Medical Research Council (MC_UU_1201/1).

